# Hand-reared wolves show attachment comparable to dogs and use human caregiver as a social buffer in the Strange Situation Test

**DOI:** 10.1101/2020.02.17.952663

**Authors:** Christina Hansen Wheat, Linn Larsson, Patricia Berner, Hans Temrin

## Abstract

Domesticated animals are generally assumed to display increased sociability towards humans compared to their wild ancestors. Dogs (*Canis familiaris*) have a remarkable ability to form social relationships with humans, including lasting attachment, a social bond based on emotional dependency. Since it has specifically been suggested that the ability to form attachment with humans evolved post-domestication in dogs, attempts to quantify attachment in wolves (*Canis lupus*), the ancestor of dogs, have subsequently been performed. However, while these rare wolf studies do highlight the potential for wolves to express attachment behaviour towards humans, the varied, and in some cases, contrasting results also emphasize the need for further testing of wolves. Here we used the standardized Strange Situation Test to investigate attachment behaviour expressed in wolves and dogs hand-reared and socialized under identical conditions. We found that 23 weeks old wolves and dogs equally discriminated between a stranger and a familiar person, and expressed similar attachment behaviours toward a familiar person. Additionally, wolves, but not dogs, expressed significantly elevated stress behaviour during the test, but this stress response was buffered by the presence of a familiar person. Wolves also expressed quantifiable fear responses to the stranger, whereas no such response was detectable in dogs. Together, our results suggest that wolves can show attachment toward humans comparable to that of dogs. Importantly, our findings demonstrate that the ability to form attachment with humans exists in relatives of the wild ancestor of dogs, thus refuting claims that such attachment is unique to post-domestication dog lineages.

## Introduction

Domestication, the evolutionary process in which animals are selected to live in human controlled environments (Price 2002), has dramatically impacted the behavioural expression in animals (Belyaev et al. 1985; Künzl and Sachser 1999; Trut 1999; Himmler et al. 2013). For instance, increased expression of human-directed sociability is considered to be more pronounced in domesticated animals when compared to their wild ancestors (Belyaev 1979; Hare, Wobber, and Wrangham 2012). The domestic dog (*Canis familiaris*) is probably the domesticated animal that has most successfully adapted to life in a human-controlled world (Wynne 2021) and many dogs live in very intimate social relationships with their owners as pets (Udell, Dorey, and Wynne 2010; Wynne 2021). Reflecting the uniqueness of this human-canine relationship, it has been suggested that dogs can form attachment bonds to their human caregivers (Topál et al. 2005; Á Miklósi and Topal 2013), and that this ability evolved post-domestication as a trait unique to dogs (Topál et al. 2005). However, whether behavioural traits seen in present day dogs are a novel product of domestication, or a target of domestication that existed as variation within pre-domestication wolf populations remains an open question. Such clarifications have significant ramifications upon our understanding of how dog domestication might have proceeded.

Attachment is a social bond based on emotional dependency formed between two individuals that endures over time (Ainsworth and Bell 1970). Originally described as the bond between a human infant and its mother, attachment behaviour constitutes any type of behaviour performed by an emotionally dependent individual to promote and maintain proximity or contact to the individual of attachment (Bowlby 1958; Ainsworth and Bell 1970). Ainsworth’s Strange Situation Procedure (SSP) is a highly influential method developed to empirically investigate the attachment bond between human infants and their primary caretaker (Ainsworth and Bell 1970). Based on the assumption that the attachment system is only activated in challenging situations (Rehn et al. 2013; Prato Previde and Valsecchi 2014), the SSP examines an infant’s attachment behaviour toward its primary caregiver under standardized test conditions of interchanging low and high emotional stress situations, which includes separation, reunion and the presence of a stranger (Ainsworth and Bell 1970). In the SSP, attachment is quantified and based on the infants expression of i) safe haven effects; which is expressed through proximity maintenance and contact seeking behaviours where the infant actively seeks to maintain physical contact with the attachment figure, and ii) secure base effects; which can be expressed as the infant’s increased willingness to engage in exploratory and/or and play behaviour when the attachment figure is present (Ainsworth and Bell 1970). Thus, attachment quantification is based on the infant’s ability to discriminate between a primary caretaker and a stranger during the SSP test conditions (Ainsworth and Bell 1970; Topal et al. 1998; Rehn et al. 2013).

Adaptation of the SSP to dogs (i.e. the Strange Situation Test (SST)) was first performed in a study just over 20 year ago (Topal et al. 1998), wherein the authors suggested that the human-dog bond is comparable to that of a parent-child attachment bond. Since then multiple studies (Gácsi et al. 2001; Valsecchi et al. 2010; Mariti et al. 2013) have used the SST to confirm that dogs express more affiliative behaviours towards their human caregiver, engage in more explorative behaviours in the presence of their human caregiver and express distress behaviour upon separation from their human caregiver when compared to a stranger. In 2005, Topál et al. were also the first to compare attachment in dogs and hand-raised wolves using the SST, finding that wolves did not discriminate between a familiar person and a stranger at 16 weeks of age. The authors suggested that an absence of a direct functional relationship between puppy-mother attachment in wild wolves could explain the wolves’ inability to form attachment bonds to their human caregivers and concluded that dogs have evolved a unique ability to show attachment toward humans post-domestication.

A total of four studies have subsequently investigated attachment behaviour using hand-raised wolves (Gácsi et al. 2005; Hall et al. 2015; Ujfalussy et al. 2017; Lenkei et al. 2020). The results from these studies vary considerably in their findings of attachment. Gácsi et al. (2005) reported that wolves up to five weeks of age expressed aggression and avoidance behaviour towards their caregiver; Ujfalussy et al. (2017) reported affiliative behaviours expressed towards a familiar person at six, 12 and 24 weeks of age with discrimination between familiar and strange people; and Lenkei et al. (2020) reported that adult wolves expressed increased contact seeking behaviour and secure base effects in the presence of a familiar person. The remaining study of the four, Hall et al. (2015), was the only one to use the SST to quantify attachment behaviour in wolves and showed that hand-raised wolf puppies up to the age of eight weeks expressed attachment in the form of proximity and contact seeking behaviour toward human caregiver. While the majority of these studies indeed highlight the potential for wolves to express attachment behaviour towards humans, the varied, and in some cases, contrasting, results also illustrate the need for further testing of wolves. First, several different tests and metrics have been used to quantify attachment or social affiliation with humans across these studies, making comparisons difficult. Second, the effort necessary to hand-raise, socialize and test wolves and dogs combined with limited animal availability highlights a fundamental challenge in domestication research on canids, namely inherently small sample sizes rarely exceeding N = 11 for wolves and N = 13 for dogs (Miklósi et al. 2003; Topál et al. 2005; Gácsi et al. 2005; Udell et al. 2008; Moretti et al. 2015; F Range et al. 2015; Marshall-Pescini et al. 2017). However, as an additional caveat to these challenges comes the high demand for conducting hand-raising, socialization and testing under identical conditions in order to decrease the risk that even subtle environmental bias affect results when working with such small sample sizes. Therefore, more efforts standardizing and replicating studies on wolves and dogs are essential in furthering our understanding of the behavioural consequences of domestication.

Here we aim to further contribute to the understanding of the evolution of the human-canine relationship. We will do so by subjecting 23 week-old wolves and dogs hand-raised and socialized under identical, standardized conditions (Klinghammer and Goodman 1987; Udell et al. 2008; Range and Virányi 2011) to the SST adapted to canids (Topal et al. 1998; Topál et al. 2005). Specifically, we will quantify safe haven and secure base effects, such as greeting and exploration respectively, as is traditionally done throughout the SST (Topal et al. 1998; Topál et al. 2005) to assess attachment in wolves and dogs. However, we will also include a separate, simultaneous quantification of basic stress and fear behaviours throughout the SST to gain novel insight on how wolves and dogs are affected by the test situation across episodes.

## Materials and Methods

### Study animals

Twelve Alaskan huskies and 10 European grey wolves were included in this study. The dogs, four females and eight males, came from two unrelated litters. The wolves, five females and five males, came from three different litters where of two litters were full siblings and the third was unrelated to the first two litters. To minimize environmental bias, including maternal effects, which are well-documented to affect the development of behavioural patterns (Clark and Galef 1982; Wilsson and Sundgren 1998; Bray et al. 2017), puppies were hand-raised and extensively socialized under standardized conditions from the age of 10 days. Each dog and wolf litter were raised separately and socialized with 24-hour presence of human caregivers for the first two months. Caregiver presence was decreased with a few hours a day from the puppies were two months. A subsequent gradual decrease of caregiver presence followed and at four months of age the puppies would spend every other night without a caregiver present. Puppies were reared in identical indoor rooms until they were five weeks old and here after given access to smaller roofed outdoor enclosures. At six weeks of age, after a week of habituation to the roofed outdoor enclosure, puppies were given access to a larger fenced grass enclosure. From the age of six weeks the puppies had free access to all three enclosures during the day and access to the indoor room and the roofed enclosure during the night. At three months of age puppies were moved to large outdoor enclosures (two enclosures of 2,000 square meters each for the wolves and dogs), in which they remained for the rest of the study period. Dogs and wolves were kept separate throughout the entire period. Behavioural observations were initiated immediately at 10 days of age and behavioural testing was initiated at six weeks of age. Caregiving, handling and socialization procedures, enrichment, testing procedures and exposure to the new environments were standardized across all three years. Hand-raisers were both male and female. Puppies were never disciplined or trained. From the age of eight weeks puppies were gradually introduced to strangers through the fence, always with the support of one or more caregivers, and were at the time of the SST accustomed to groups of up to 10-15 strangers.

### Experimental design

All wolves and dogs were tested in the SST at the age of 23 weeks (dogs: 23 weeks ± 0; wolves: 23 weeks ± 0.3). The experimental design was identical to that of Topál et al. (2005). Briefly, the SST adapted to dogs consists of seven experimental episodes, each lasting two minutes, in which the presence and absence of a familiar person and a stranger in a test room with the focal animal is alternated (Table 1).

**Table 1.** Strange Situation Test procedure. In the seven episodes in the Strange Situation Test a familiar person (F) and/or a stranger (S) is present in the test room with the focal animal (except for episode 5 where the animal is alone). Each episode is structured differently. The procedure is identical to the study of Topál et al. (2005).

| Episode | Present | Minutes | Structure of episode |
| --- | --- | --- | --- |
| 1 | F | 0-2 | F leads the animal into the test room, closes the door and sits down in one of two chairs and reads from a paper in silence. After 1 min F initiates play with the animal. F stops playing after 2 min as S enters the room. |
| 2 | F+S | 2-4 | S enters the room and stops for up to 5 s, allowing the animal to greet, and then sits down in the vacant chair. After 30s S initiates a friendly chat with F. After 30s S stops chatting with F, stands up and initiates play with the animal. F then leaves as quite as possible. |
| 3 | S | 4-6 | S continues to play/initiates play with the animal. After 1 min S stops playing and returns to the chair. If the animal initiates contact, S is allowed to reciprocate physical contact by petting it. |
| 4 | F | 6-8 | F calls the animal from outside the room. After entering the room, F stops for up to 5s to allow the animal to greet and then goes to the chair and sits down. S leaves. F initiates play with the animal for 1 min and then returns to the chair. If the animal initiates contact, F is allowed to reciprocate physical contact by petting it. At the end of the episode F says 'I must go, stay here' and leaves the room. |
| 5 | - | 8-10 | The animal is alone in the room. |
| 6 | S | 10-12 | S enters the room and stops for up to 5s to allow the animal to greet, and then initiates play with the animal. After 1 min S sits down in the chair. If the animal initiates contact, S is allowed to reciprocate physical contact by petting it. |
| 7 | F | 12-14 | F calls the animal from outside the room. After entering the room, F stops for up to 5s to allow the animal to greet. S leaves the room while F stimulates the animal to play for 1 min and then sits down in the chair. If the animal initiates contact, F is allowed to reciprocate physical contact by petting it. |

The familiar person was a primary, female caregiver, who had raised all the litters from 10 days of age and was the hand-raiser who had spent most time with the animals. The female stranger had never met the dogs or the wolves prior to the experiment. The same familiar person and stranger were used in all tests. In the 6×6 meter test room, which was familiar to both dogs and wolves, two chairs were placed in the middle of the room, 2 meters from each other and facing in the same direction. Seven toys from the animal’s home enclosure, such as balls, rope and rubber toys, were distributed across the floor in the test room. Familiar toys were used to avoid the risk of eliciting a neophobic response. Tests were filmed with two diagonally mounted GoPro cameras (model Hero, 3, 3+, 4, GoPro Inc.).

### Behavioural scoring - SST

Following the procedures in Topál et al. (2005) a total of seven behaviours were quantified using an ethogram (Table 2). These seven behaviours included 1) greeting, following, physical contact and standing by the door – all categorized as *safe haven effects*, which are expressed as a mean to maintain proximity or physical contact with the attachment figure (Ainsworth and Bell 1970); 2) exploration and play – both categorized as *secure base effects*, which can be expressed more in the presence of the attachment figure (Ainsworth and Bell 1970), and 3) passive behaviour – categorized as *other behaviour* related to other aspects of the social and physical environment (Topál et al. 2005). Behaviours were further divided into a) continuously measured behaviours, which included exploration, passive behaviour, physical contact, social play and standing by the door, and b) scored behaviours, which included following and greeting (Table 2, Table S1).

**Table 2.** Ethogram, SST. Behavioural categories coded following Topál et al. (2005), including a) Continuously measured behaviours and b) Scored behaviours. All continuous behaviours were scored as both frequency and duration.

| Behaviour | Definition |
| --- | --- |
| *a) Continuously measured behaviours* |  |
| **Exploration** | Activity directed towards non-movable aspects of the test room (not including toys), including sniffing, distal visual inspection (staring or scanning), close visual inspection or oral examination, while F and/or S are present and during episode 5 when the animal is alone. |
| **Passive behaviour** | Sitting, standing or lying down without any orientation towards the environment (this includes grooming), while F and/or S are present, and during episode 5 when the animal is alone. |
| **Physical contact** | Bodily contact initiated by F or S (e.g. petting, touching) or the animal. |
| **Social play** | Motor activity performed when interacting with F or S; including running, jumping, active physical contact and chasing toys. |
| **Stand by the door** | Standing within 1 meter of the door and facing towards the door. |
| *b) Scored behaviours* |  |
| **Following** | Conditional scoring between 0 and 3 of following F and S leaving the room while the other person stays behind. **Score 0:** no orientation towards the leaving person at all, or only for less than 1s. **Score 1:** orientation towards the leaving person for more than 1 s. **Score 2:** following the leaving person to the door. **Score 3:** trying to get through the door or standing by the door for more than 1s. The mean based on scores from episode 3, 4 and 7 is used as total score. |
| **Greeting** | The behaviour of the animal towards the entering F or S, scored using one of five categories: **approach initiation (+1):** the animal moves towards the entering person; **full approach (+1):** the animal approaches the entering person until physical contact is made; **avoidance (-1):** avoidance behaviour towards the entering person, such as backing or getting out of the way of the entering person; **durable physical contact upon greeting (+1/2):** the animal spends more than 3s in bodily contact with the entering person; **delay of approach (-1/2):** when F or S enters, the animal hesitates to initiate any form of approach for more than 5s. Scores are summed up to a total score (maximum 5 since each person entered twice) |

*F = Familiar person, S = Stranger.*

### Behavioural scoring – stress and fear behaviours

As a compliment to the SST we also scored various behaviours previously used to quantify stress and fear in wolves and dogs (Ujfalussy et al. 2017; Cimarelli et al. 2021) throughout the test using a separate ethogram (Table 3). Originally the behaviours crouching, growling, piloerection, pacing, retreating, startle, tail tuck, and yawning were included in this ethogram. However, with the exception of pacing, crouching and tail tuck, all other behaviours occurred so rarely, or not at all, that we chose to exclude these behaviours from further consideration (but see Table S2 for full quantification of stress and fear behaviours). Pacing, crouching and tail tuck were all quantified as state behaviours using both duration and frequency for extraction of fractions and were not necessarily mutually exclusive.

**Table 3.** Ethogram, stress and fear behaviours. Behavioural categories coded for stress and fear behaviours occurring as state and events during the SST.

| Behaviour | Definition |
| --- | --- |
| **Crouching** | Lowered body position in which the back is curved. Can be accompanied by tucking of the tail. |
| **Pacing** | Walking or trotting at a steady speed without any exploratory purpose or obvious focus on the surroundings. |
| **Tail tuck** | The tail is tucked down between the hind legs, the tail might touch the underside of the stomach |

Behavioural scoring for both attachment, and stress and fear behaviours was carried out using the software program BORIS v. 2.97 (Friard and Gamba 2016). For the SST of the recorded tests, 25% were independently coded by two of the authors. Cronbach’s Alpha was calculated and inter-observer reliability was high for all continuous behaviours (exploration: 0.986; passive behaviour: 0.978; social play: 0.985; stand by door: 0.989; physical contact: 0.987).

### Statistical analyses

For all statistical analyses we used the software SPSS Statistics v. 25.

To account for slight variations in durations of episodes across tests (because of the time it took for the test persons to enter and exit the room), we used the relative proportion of the time spent on each behaviour for every episode for all individuals. Upon testing for normality using a Shapiro-Wilk test (Razali and Wah 2011), we found that the two variables passive behaviour and social play were not normally distributed. We therefore arcsine transformed these two variables prior to statistical testing. Arcsine transformation is commonly used for proportional data (McKillup 2012; Cohen et al. 1983). For one wolf the test was aborted prematurely and as a result episode 6 and 7 were excluded for this individual, i.e. N=9 for the wolves in some comparisons. We present the mean and SE for the untransformed data in our figures (McDonald 2009, Table S2).

The behaviours Greeting, Following, Physical contact, Standing by the door, Exploration, Social play and Passive behaviour were divided into two main categories: 1) ‘In the presence of the familiar person’, which refers to those episodes in which the familiar person was present (1, 2, 4, 7), and 2) ‘In the presence of the stranger’, which refers to those episodes where the stranger was present (2, 3, 6). For the three behaviours Pacing, Crouching and Tail Tuck, the results are given for each of the seven episodes. We used a General linear model (GLM) for repeated measurements where the proportions of the presence of the familiar person and the stranger were classified as the within-subject factor, and dogs and wolves as the between-subject factor. The least significant difference (LSD) pairwise multiple comparison test, i.e. multiple individual t-tests between all pairs of groups, was used for the results in Table 4.

**Table 4.** Posthoc comparisons of episodes for stress and fear behaviours. The least significant difference (LSD) pairwise multiple comparison test for the relevant stress and fear behaviours in wolves. Given for each behaviour is Episode, Episode comparison (comp.), mean difference (diff.), Standard (Std.) error, p-value (p). and 95% confidence intervals (Low and Up). Significant p-values are given in bold italic.

| Episode | Episode comp. | Mean diff. | Std. Error | p | 95 CI Low | 95 CI Up |
| --- | --- | --- | --- | --- | --- | --- |
| *a) PACING* |  |  |  |  |  |  |
| **4** | 1 | -1.015 | 0.461 | 0.059 | -2.078 | 0.048 |
|  | 2 | -1.204 | 0.214 | **>0.001** | -1.696 | -0.711 |
|  | 3 | -1.507 | 0.291 | **0.001** | -2.179 | -0.835 |
|  | 5 | -0.943 | 0.277 | **0.009** | -1.583 | -0.303 |
|  | 6 | -1.843 | 0.436 | **0.003** | -2.848 | -0.838 |
|  | 7 | 0.868 | 0.262 | **0.011** | 0.263 | 1.473 |
| **7** | 1 | -1.883 | 0.343 | **0.001** | -2.675 | -1.091 |
|  | 2 | -2.072 | 0.328 | **>0.001** | -2.828 | -1.315 |
|  | 3 | -2.375 | 0.209 | **>0.001** | -2.856 | -1.894 |
|  | 5 | -1.811 | 0.309 | **>0.001** | -2.523 | -1.1 |
|  | 6 | -2.711 | 0.555 | **0.001** | -3.991 | -1.431 |
| *b) CROUCHING* |  |  |  |  |  |  |
| **2** | 1 | 5.860 | 2.386 | **0.04** | 0.359 | 11.361 |
|  | 3 | 5.173 | 2.192 | **0.046** | 0.118 | 10.228 |
|  | 4 | 5.860 | 2.386 | **0.04** | 0.359 | 11.361 |
|  | 5 | 5.860 | 2.386 | **0.04** | 0.359 | 11.361 |
|  | 6 | 4.013 | 2.167 | 0.101 | -0.984 | 9.01 |
|  | 7 | 5.860 | 2.386 | **0.04** | 0.359 | 11.361 |
| **6** | 1 | 1.847 | 0.801 | 0.05 | 0.001 | 3.693 |
|  | 3 | 1.16 | 0.635 | 0.105 | -0.304 | 2.625 |
|  | 4 | 1.847 | 0.801 | 0.05 | 0.001 | 3.693 |
|  | 5 | 1.847 | 0.801 | 0.05 | 0.001 | 3.693 |
|  | 7 | 1.847 | 0.801 | 0.05 | 0.001 | 3.693 |
| *c) TAIL TUCK* |  |  |  |  |  |  |
| **2** | 1 | 2.206 | 1.224 | 0.109 | -0.617 | 5.029 |
|  | 3 | 1.927 | 1.291 | 0.174 | -1.05 | 4.904 |
|  | 4 | 2.246 | 1.196 | 0.097 | -0.512 | 5.003 |
|  | 5 | 2.303 | 1.181 | 0.087 | -0.421 | 5.027 |
|  | 6 | 0.632 | 1.457 | 0.676 | -2.727 | 3.992 |
|  | 7 | 2.842 | 1.178 | **0.042** | 0.125 | 5.559 |
| **6** | 1 | 1.574 | 0.65 | **0.042** | 0.074 | 3.074 |
|  | 3 | 1.295 | 0.497 | **0.031** | 0.149 | 2.441 |
|  | 4 | 1.613 | 0.614 | **0.03** | 0.197 | 3.03 |
|  | 5 | 1.670 | 0.648 | **0.033** | 0.176 | 3.164 |
|  | 7 | 2.21 | 1.068 | 0.072 | -0.254 | 4.674 |

### Ethical statement

Daily care and all experiments were performed in accordance with guidelines and regulations under national Swedish Law. The experimental protocols in this study were approved by the Ethical Committee in Uppsala, Sweden (approval number: C72/14). Facilities and daily care routines were approved by the Swedish National Board of Agriculture (approval number: 5.2.18-12309/13). As required by national law in Sweden, all hand-raisers working with the puppies were ethically certified and trained to handle animals.

## Results

### 1) Attachment behaviours - safe haven effects

#### Greeting

Greeting was only scored when the familiar person or the stranger entered the room, which occurred during episode 2, 4, 6 and 7. Greeting was expressed significantly more towards the familiar person than towards the stranger (*F*1,19=10.225, *P*=0.005, N_wolf_ =9, N_dog_ =12, Figure 1a, Table S3). There was no difference between dogs and wolves in their expressions of greeting behaviour (*F*1,19=0.637, *P*=0.435, Figure 1a, Table S3) and no interaction (*F*1,19=2.113, *P*=0.162, Figure 1a, Table S3).

**Figure 1.**
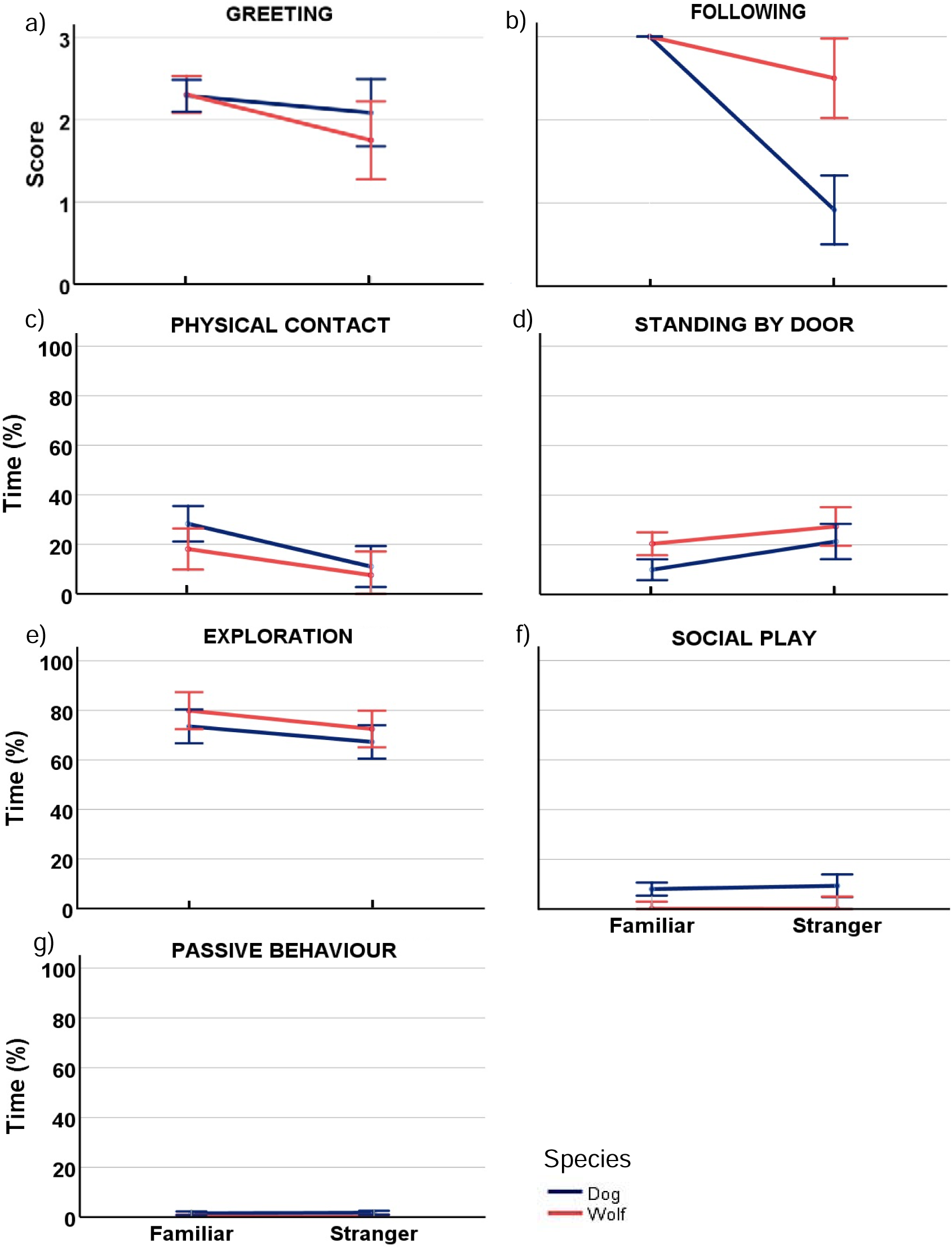
Attachment behaviours, discrimination between familiar person and stranger. Mean score for dogs (blue) and wolves (orange) for a) Greeting and b) Following a familiar person and a stranger, mean proportion of time dogs and wolves spent on c) Physical contact, d) Standing by the door, e) Exploration, f) Social play and g) Passive behaviour, in the presence of a familiar person and a stranger. Error bars represent 95% confidence intervals. See Table S3 for all statistical outputs.

#### Following

The familiar person was significantly more likely to be followed when leaving the room compared to when the stranger left the room (*F*_1,19_=73.134, *P*<0.001, Figure 1b, Table S3) and there was a species difference in the proportion of following (*F*_1,19_=27.473, *P*<0.001, Figure 1b, Table S3) Both wolves (paired t-test: *t*_9_=2.683, *P*=0.028, Table S4) and dogs (*t*_12_=9.449, *P*<0.001, Table S4) followed the familiar person more than the stranger. There was a significant interaction effect between species and test person (*F*1,19=27.473, *P*<0.001, Figure 1b, Table S3), suggesting that the difference between familiar person and stranger was greater in dogs than in wolves.

#### Physical contact

Physical contact with the familiar person was significantly more common than with the stranger (*F*_1,20_=12.223, *P*=0.002, N_wolf_ =9, N_dog_ =12, Figure 1c, Table S3). Wolves and dogs did not differ in their expression of physical contact (*F*_1,20_=2.914, *P*=0.104, Figure 1c, Table S3) and there was no interaction (*F*_1,20_=0.717, *P*=0.408, Figure 1c, Table S3).

#### Standing by the door

Both wolves and dogs stood more by the door when the stranger was in the room and the familiar person was absent, than when the familiar person was in the room (*F*_1,20_=18.346, *P*<0.001, Figure 1d, Table S3). Wolves stood more by the door compared to dogs (*F*_1,20_=5.391, *P*=0.031, Figure 1d, Table S3), but there was no interaction (F_1,20_=1.050, *P*=0.318, Figure 1d, Table S3).

### 2) Attachment behaviours - secure base effects

#### Exploration

Exploration was significantly more common in the presence of the familiar person than in the presence of the stranger (*F*_1,20_=7.968, *P*=0.011, N_wolf_=10, N_dog_=12, Figure 1e, Table S3). There was no difference between wolves and dogs in the expression of explorative behaviour (*F*_1,20_=1.928, *P*=0.180, Figure 1e, Table S3) and there was no interaction (*F*_1,20_=0.056, *P*=0.815, Figure 1e, Table S3).

#### Social play

The expression of social play was not affected differently by the presence of the familiar person and the stranger (*F*_1,20_=0.371, *P*=0.549, N_wolf_ =10, N_dog_ =12, Figure 1f, Table S3). Dogs were significantly more playful than wolves (*F*_1,20_=12.761, *P*=0.002, Figure 1f, Table S3), but there was no interaction (F_1,20_=0.4443, *P*=0.513, Figure 1f, Table S3).

### 3) Other behaviours

#### Passive behaviour

The expression of passive behaviour was low in both wolves and dogs and not affected differently by the presence of the familiar person and the stranger in either species (*F*_1,20_=0.053, *P*=0.821, N_wolf_ =10, N_dog_ =12, Figure 1g, Table S3). Dogs were significantly more passive than wolves (*F*_1,20_=14.396, *P*=0.001, Figure 1g, Table S3), but there was no interaction (*F*_1,20_=0.112, *P*=0.741, Figure 1g, Table S3).

### 4) Stress behaviours

#### Pacing

There was a significant difference in the expression of pacing between the seven episodes (F=8.528, p<0.001, df=6, Figure 2a, Table S5) and there was a significant difference between dogs and wolves (F=48.101, p<0.001, df=1, Figure 2a, Table S5). There was no difference in pacing between the seven episodes in dogs (F=1.645, p=0.149, df=6, Figure 2a: Table S6). In wolves, there was an overall difference in pacing between the seven episodes (F=10.449, p<0.001, df=6, Figure 2a, Table S7). Compared to all other episodes, wolves were pacing significantly less in episode 4 and 7 when they were reunited with the familiar person after having been alone with the stranger (Figure 2a, Table 4a, see Table S7 for all posthoc comparisons). The only exception was episode 1, in which the wolves had just entered the test room alone with the familiar person, and episode 4. Here the difference in pacing was not significant (p = 0.059, Figure 2a, Table 4, Table S7).

**Figure 2.**
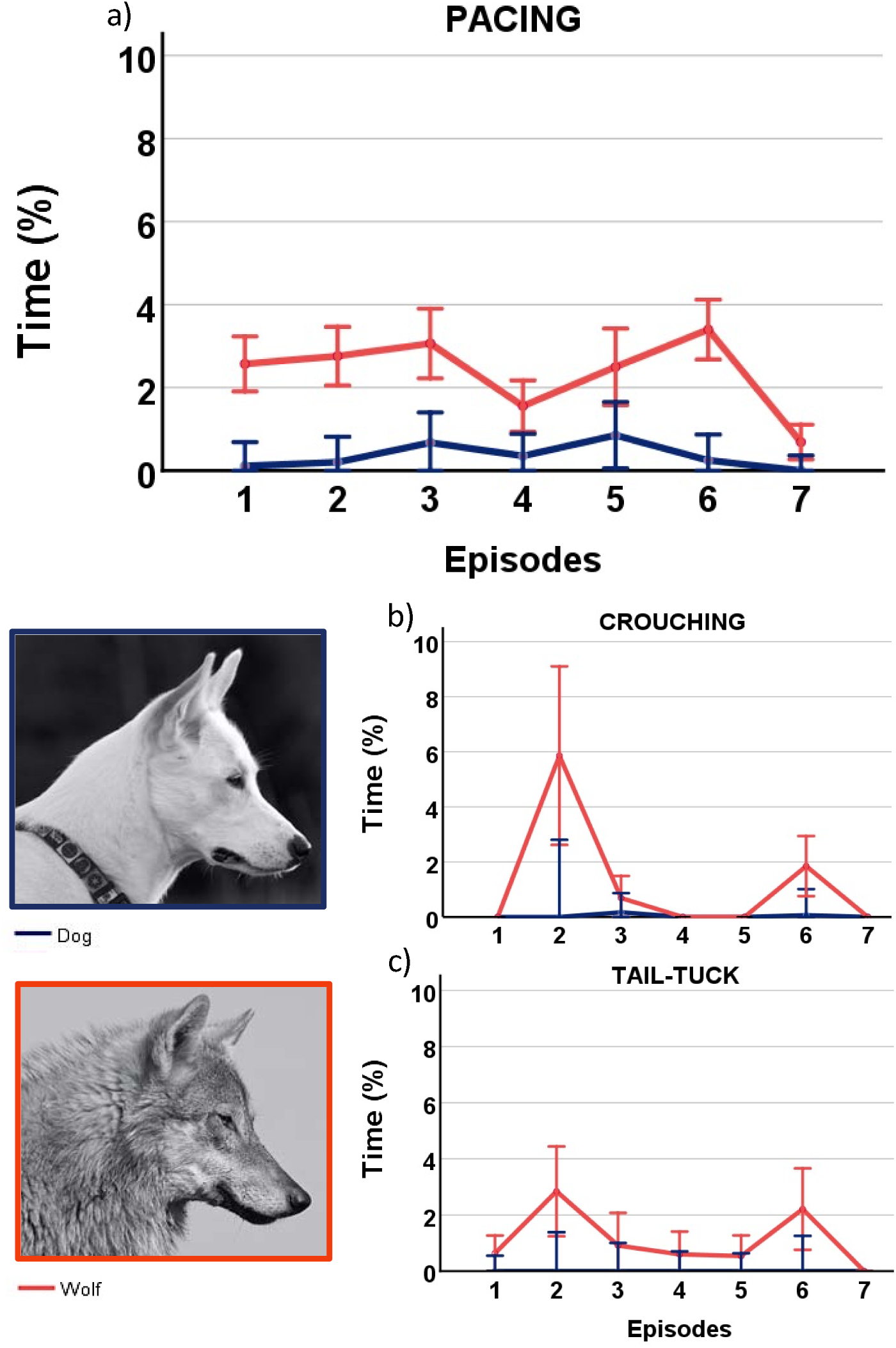
Stress and fear behaviours during the SST. Mean proportion of a) Pacing, b) Crouching and c) Tail tucking occurring in each of the seven episodes in the SST in dogs (blue) and wolves (orange). Error bars represent 95% confidence intervals. See Tables S5-S10 for all statistical outputs.

### 5) Fear behaviours

#### Crouching

There was a significant difference in the expression of crouching between the seven episodes (F=7.256, p<0.001, df=6, Figure 2b, Table S5) and there was a difference between dogs and wolves (F=9.005, p=0.007, df=1, Table S5). There was no difference in crouching between the seven episodes in dogs (F=0.882, p=0.513, df=6, Figure 2b, Table S8). In wolves, there was an overall difference in crouching between the seven episodes (F=5.414, p<0.001, df=6, Figure 2b, Table S9). Crouching behaviour was only expressed in wolves in episodes in which the stranger was present. Compared to all other episodes, wolves were crouching significantly more in episode 2 when the stranger entered the test room for the first time (Figure 2b, Table 4b, see Table S9 for all posthoc comparisons of episodes). The only exception was episode 6, in which stranger enters the room for the second time (p = 0.101, Table 4b, Table S9). There was also a noticeable peak in crouching behaviour in episode 6 compared to all other episodes, except for episode 2, but while the majority of p-values were just at the 0.05 level and a few above, none were significant (Figure 4b, Table 4b, Table S9).

#### Tail tuck

There was a significant difference in the expression of tail-tuck between the seven episodes (F=4.210, p=0.001, df=6, Figure 2c, Table S5) and there was a difference between dogs and wolves (F=5.542, p=0.029, df=1, Figure 2c, Table S5). This behaviour was exclusively expressed in wolves and there was an overall difference of tail-tuck between the seven episodes (F=3.102, p=0.012, df=6, Figure 2c, Table S10). There was a peak in the expression of tail tucking in the episodes when the stranger entered the room (episode 2 and 6). However, while tail tucking was significantly more expressed in episode 6 compared to any other episode, except for episode 2 and 7 (Figure 4c and Table 4c, see Table S10 for all posthoc comparisons of episodes). Tail tucking in episode 2 was significantly higher than in episode 7, which is the episode when the familiar person re-entered the room (Figure 4c, Table 4c, Table S10).

## Discussion

Here we demonstrate that 23 weeks old wolves and dogs equally discriminate between a stranger and a familiar person, and show similar attachment behaviours toward a familiar person through the expression of safe haven and secure base effects. Additionally, while wolves, but not dogs, expressed significantly elevated stress behaviour during the test in the form of pacing, this stress response was buffered by the presence of a familiar person. Wolves also expressed quantifiable fear responses to the stranger, whereas no such response was detectable in dogs. Together, our results suggest that wolves can show attachment toward humans comparable to that of dogs. Importantly, our findings demonstrate that the ability to form attachment with humans exists in relatives of the wild ancestor of dogs, thus refuting claims that such attachment is unique to post-domestication dog lineages.

Our results thus represent a stark contrast to an earlier study by Topal et al. (2005), who upon comparing 16-week-old wolves and dogs found that wolves did not discriminate in their attachment behaviour toward a human caregiver and a stranger while dogs did discriminate in favour of the caregiver. The authors suggested that wolves did not have the need for a strong bond with their mother in the wild after 6-8 weeks of age, and this could explain their lack of attachment towards a human caregiver at the age of 16 weeks. Yet here we demonstrate that wolves aged 23 weeks of age are capable of discriminating between a stranger and a familiar person and showing attachment to a human caregiver. Our results are further supported by two previous studies documenting that wolves aged up to 24 months (Ujfalussy et al. 2017) and wolves older than 1.5 years of age (Lenkei et al. 2020) also express attachment behaviours toward their human caregivers in other test set-ups than the SST. Furthermore, it is important to note that the wolves in the study by Topal et al. (2005) had been relocated to an animal park up to two months before the SST test was conducted (Hall et al. 2015; Virányi et al. 2008). Thus, at the time of testing, dogs were still living with their caregivers, but wolves were not, i.e. wolves and dogs were not kept under similar conditions prior to testing. Because environmental factors significantly affect behavioural development (Zimen 1987) and experimental outcomes in studies comparing wolves and dogs (B Hare 2002; Udell, Dorey, and Wynne 2008), this environmental difference between the test animals in the study by Topal et al. (2005) and our study could be one explanation for the contrasting results.

The attachment system toward the familiar person was activated in our 23 week old wolves during the SST, and the wolves expressed attachment behaviour comparable to those reported in adult dogs (Topal et al. 1998; Gácsi et al. 2001; Prato-Previde et al. 2003), chimpanzees (van IJzendoorn et al. 2009) and human infants (Ainsworth and Bell 1970) when subjected to the SST or SSP. Specifically, our wolves expressed safe haven and secure base effects similar to that of identically raised dogs towards, or in the presence of, the familiar person, but not the stranger, which included significantly pronounced contact seeking, proximity maintenance and exploration. The expression of passive behaviour and social play was limited in both wolves and dogs. Though dogs expressed higher levels of both those behaviours compared to wolves, the expression of neither passive behaviour nor social play was dependent on the presence of the familiar person or the stranger in either species. We note that the limited occurrence of passive behaviour and social play during the test might have impaired our ability to adequately detect a difference in these behavioural expressions in the presence of a familiar person compared to a stranger (Rehn et al. 2013). Additionally, wolves engaging in play with a human are likely rare (Hansen Wheat and Temrin 2020) and wolf hybrids show significantly decreased expression of human-directed play behaviour when compared with dogs (Hansen Wheat et al. 2018). Therefore, the wolves’ limited engagement in social play during the SST is likely based on a general lack of interest in human-directed play, and does not necessarily represent an expression of low attachment (i.e., secure base effect) to the familiar person.

In dogs, acute stress responses to solitary, restricted confinement can manifest as repetitive movement patterns such as pacing (Beerda et al. 1997). While the test situation in the SST is not strictly solitary *per se*, as humans are present for the majority of the test, the dogs and wolves were separated from their littermates for the purpose of the test and confined to the restricted space of the test room. The wolves had an exaggerated stress response to the test situation compared to the dogs and expressed significantly increased pacing behaviour throughout the test. However, in the two reunion episodes in which the familiar person re-enters the room after the wolves had been alone with the stranger, pacing was significantly reduced. This notable reduction in stress response suggests that the familiar person acted as a social buffer for the wolves in an aversive situation (Hennessy et al. 2009). The facilitation of comforting effects in stressful situations by familiar conspecifics is well-known in various species (von Holst 1998; Hennessy et al. 2006) and has recently been demonstrated among captive wolf pack members (Cimarelli et al. 2021). Social buffering is believed to be related, though not identical to, attachment and secure base effects (Lenkei et al. 2020), and the social buffering demonstrated in our study thereby strengthens our main conclusion that wolves are capable of showing attachment to a human caregiver.

Furthermore, wolves, but not dogs, had a quantifiable fear response to the strange person entering the test room, expressed as pronounced crouching and tail tucking. This is in line with previous findings of strangers, but not familiar humans, eliciting crouching and tail tucking in hand-raised wolves (Ujfalussy et al. 2017), and further lends support to the general assumption that hand-raised wolves do not generalize their socialization to strangers (Zimen 1987; Klinghammer and Goodman 1987) as dogs do (Udell 2015). The absence of stress and fear responses in the dogs in the test situation could thus be an effect of domestication. However, no matter the causal explanation for the test situation not eliciting quantifiable stress and fear behaviours in the dogs, the fact that this was the case in the wolves highlights a potential problem in the way that wolves and dogs are generally tested for behavioural comparisons. While the demand for testing wolves and dogs under similar conditions is crucial in order to be able to compare behavioural outcomes, such test setups inherently favor conditions which must be categorized as being more natural to dogs than to wolves. As demonstrated here, this is likely to elicit an aversive response to the test situation itself in wolves, but not dogs, and this should perhaps be taken into consideration as we go forward in our pursuit to expand our knowledge about behavioural evolution during dog domestication.

In sum, our results add to a slowly, but steady, growing body of evidence that wolves are capable of expressing attachment behaviour towards human-caregivers (Hall et al. 2015; Ujfalussy et al. 2017; Lenkei et al. 2020). Additionally, these studies highlight that wolves across a wide range of ontogenetic stages, and not just very young wolf puppies, but also adult wolves, possess this capability. Together, the collective evidence from these wolf studies suggests that the narrative that the ability to form attachment with humans is exclusive to dogs is no longer tenable.

## Acknowledgments

We wish to thank Stockholm University for funding this study, Yvonne Silfverblad who was the stranger in the SST, our hand-raisers Anna Björk, Marjut Pokela, Charles Gent, Åsa Lycke, Erika Grasser, Joanna Schinner, Yrsa Andersson, Christoffer Sernert and Mija Jansson, the staff at Tovetorp Zoological Research Station and Dr. Józef Topál for providing helpful insight during data analyses.

